# Integrating science and citizen science: the dusky grouper (*Epinephelus marginatus*) sustainable fishery of Copacabana, Rio de Janeiro, Brazil

**DOI:** 10.1101/759357

**Authors:** Alpina Begossi, Svetlana V. Salivonchyk

## Abstract

We followed landings of dusky grouper, *Epinephelus marginatus*, from 2013 to 2019. We observed 1,896 individuals of dusky grouper, *Epinephelus marginatus*, in Copacabana, Rio de Janeiro, from September 2013 to February 2019. The total weight of the catches was 6,065.57 kg, with an average of 1,442.50 kg/year and a std of 147.30 kg.

We integrated fishers in our study through citizen science (CS): individuals were trained to monitor grouper gonads and supplied information on fishing spots and prices. After comparing catch curves (based on weight) and curve prices (in the Brazilian monetary currency of reals), our results showed that catches in the Copacabana fishery have been stable (the results of the Kruskal-Wallis test showed no significant difference for either the weight of the catches or the average prices of dusky groupers in the years compared). Copacabana has been a sustainable fishery when considering its catches of dusky grouper. This is a very important result for conservation and management, considering the importance of small-scale fisheries in terms of their low fishing efforts and their possible effects on vulnerable species, as well as their ecological and economic importance in developing countries. Citizen science, alomng with local ecological knowledge, helps integrate research and fisheries as well as researchers and fishers and allows for larger sampling efforts and management training for fishers.

## Introduction

The dusky grouper, *Epinephelus marginatus*, inhabits rocky reefs and is a slow-growing, protogynous hermaphroditic, late-maturing fish that can form spawning aggregations; since 2004, it has been considered an endangered fish by the IUCN Red List [1].

### Small-scale fisheries

Small-scale fisheries (SSFs) can be sustainable; several examples of Caribbean and Latin American SSFs are given by Salas et al. [2]. In Brazil, SSFs are of overwhelming importance, providing protein for poor coastal communities and riverine inhabitants; SSFs provide approximately half of the country’s fish production, but in some regions, it can reach higher rates [3]. However, for slow-growing and late-maturing species, SSFs can have an impact on populations [4]. This study shows that, for dusky grouper caught in Copacabana, Rio de Janeiro, there is no such impact, since production has been stable for years.

It is important to recall that SSFs are defined by several attributes [3,5,6]:

a. Small-scale fisheries (SSFs) are important at the local, regional, national and global levels;
b. SSFs use less energy-intensive fishing and usually operate in inshore waters;
c. SFFs discard little or no fish;
d. SFFs employ 25 times more people, and provide a food source for millions; and
e. SFFs lack infrastructure, often occur in remote areas and have low political power.

### Citizen science, local knowledge and the Copacabana fishery

Disciplines occupy different niches, and ecology came to be a separate niche (as did human ecology) in the postwar period, altering its identity from a soft to a hard science; however, the field of ecology has never lost its links to the physical and social sciences [7]. The understanding of the complexities of fisheries management can help to re-establish the balance between the physical and social sciences and conservation ecology by demonstrating that scientific research could be aided in obtaining positive outcomes for fisheries management by other forms of knowledge.

Currently, local knowledge (LEK) is still under scrutiny with respect to its usefulness and its acceptance in biological or fisheries science [8]. Here, we argue that LEK is very important for data-poor countries; through knowledge from fishers (i.e., LEK) or the public (i.e., CS), we can reduce the costs of time and money by increasing the efficiency of ecological data collection. Tropical countries, in particular, lack the infrastructure for data collection, and many have high biodiversity with several species for which there is no knowledge at all. In Brazil, for example, among the 65 marine species most often consumed by marine small-scale fisheries, 33% are decreasing, and 54% have an unknown status [9]. In the freshwater fisheries of the Amazonian rivers, among the 90 fish species mentioned by fishers as being consumed, 78% have an unknown biological or ecological status [10].

Local, folk, ecological and traditional knowledge bases are sources of information from local people. Such information is often empirically based and rich in ecological or biological information, which is often acquired over several years or centuries (for definitions see: [11-13]). This example, given for Brazil, reflects the reality of other countries in continents such as South America and Africa: often, there is little systematic data collection, but studies of local knowledge have been used to accumulate several important data points on local species. This is the case for coastal fisheries and for the data collected for dusky grouper.

Since 1997, one of the authors has been studying the Copacabana SSF through many research projects (FAPESP 1997, 2001, 2004, 2006, 2007 and 2014). A study was conducted by Nehrer and Begossi [14] about the fishing activities in this fishery, where the main fishing techniques were set gillnet followed by hook and line; in this period, diving was still not dominant. Groupers, led by dusky grouper but also including *Mycteroperca acutirostris* [15], were one of the notable species caught [16].

The first study conducted directly on dusky grouper in Copacabana was published by Begossi and Silvano [17]; in this study, 40 individuals of dusky grouper were collected and their stomach contents and gonads were quantified. Fishers helped by providing local knowledge (e.g., diet, habitat, and spawning) and citizen science (e.g., participation in fishing trips for information on fishing spots).

In another study, fishers were active participants in a research project as citizen scientists (besides local knowledge); the fishers were trained to observe the gonads of dusky groupers, and during the 21-month study, 800 dusky groupers were observed [18]. We continued the study of the Copacabana fishery, following catches and getting information through science and citizen science with the objective of having more information and comparing yearly data to understand if catches of dusky grouper were increasing, stable or decreasing in Copacabana. This information is essential to predict the effect of fisheries and to formulate management strategies for vulnerable species.

## Materials and methods

### Study site

The “Colônia de Pescadores do Posto 6” includes a small-scale fishing community in Copacabana beach, created in 1923. Groupers have been, and are, a target fish because of their high price [14]. Fishing is performed in small motorboats through set gillnets, hooks and lines and by spear fishing [15]. Recently, spear fishing through diving has become important, especially among young fishers.

### Procedures

Fieldwork occurred from September 25, 2013, to February 11, 2019, at the landing point of the Colonia de Pescadores Z-13 (“Posto 6”), in Copacabana. The landing point includes “selling boxes” where fishermen sell the fish that is landed. Sometimes, this is not done by fishers but by middlemen who receive the landings and sell the fish.

Two fishers were trained to use the same protocol as that in a previous study [18]: measurement of the TL (total length in cm) and weight (kg) of the dusky grouper, and dissection of the fish to observe its gonads (i.e., whether or not the fish was mature, and whether or not it had visible eggs). The procedure of macroscopically observing the mature gonads of the fish was previously used in studies with snook (*Centropomus undecimalis*) [19], bluefish (*Pomatomus saltatrix*) [20] and dusky grouper [21,22]. “Macroscopic observation” means that the gonads were observed by a naked eye. Monthly visits of approximately 3-5 days each were performed to follow the study and the groupers; when possible, the Copacabana fishery was visited twice a month for fieldwork. The two fishermen trained since the previous study [18] continued to collaborate on this research; they are also “fish cleaners”, who clean and cut fish fillets.

### Comparing yearly catches

We compared grouper catches from September 2013-September 2018 (4 full years) and also from September 2013-February 2018 (6 partial years). We built two histograms of the distributions for daily and monthly catches (in kg), which showed that the size distributions of the catches are not normal. For this reason, we used the nonparametric Kruskal-Wallis test.

For the Kruskal-Wallis test, we used data of the monthly catches in kg.

Two versions of calculations were made: 1) for 4 full years and 2) for 6 years (including full and partial years).

Thus, in the first version, we had 48 observations, and in the second version, we had 54 observations.

The sum of the ranks of the total observations in the first version is expressed as 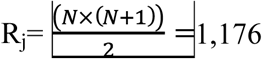; in the second version, R_j_ is 1,485. Please see the Supporting material for details.

### Comparing yearly prices

Normalized histograms were built for the grouper price distributions in the different years (2016-2018, partial years) and showed non-normal distributions.

The Kruskal-Wallis test was chosen as a nonparametric statistical test of the similarity of sample distributions when several variables (i.e., more than 2) are available. For the implementation of this test, it is preferable not to have a very long sequence of data (i.e., less than 60 samples) and to have samples with no more than two times the difference in length.

Considering the aforementioned, for our analysis, we used prices averaged per day and divided the 2017 data into quarters. Thus, we obtained 5 independent samples with a similar number of observations corresponding to time periods of approximately the same length. The total number of observations was n=128.

The sum of the ranks of the total observations was 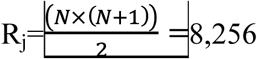. For details, consult the Supporting material.

## Results

We observed 1,896 individuals of dusky grouper, *Epinephelus marginatus*, in Copacabana, Rio de Janeiro, from September 2013 to February 2019. The total weight of the catches was 6,065.57 kg, with an average of 1,442.50 kg/year and a std of 147.30 kg. The weight, per month, of the full years of sampling (i.e., 2014-2017) is shown in Fig 1. Our sample from landings of dusky grouper in Copacabana showed an average weight of 3.20 kg (n=1,896; std=2.74) and length of 54.47 cm (n=1,544; std=12.11).

**Fig. 1.**
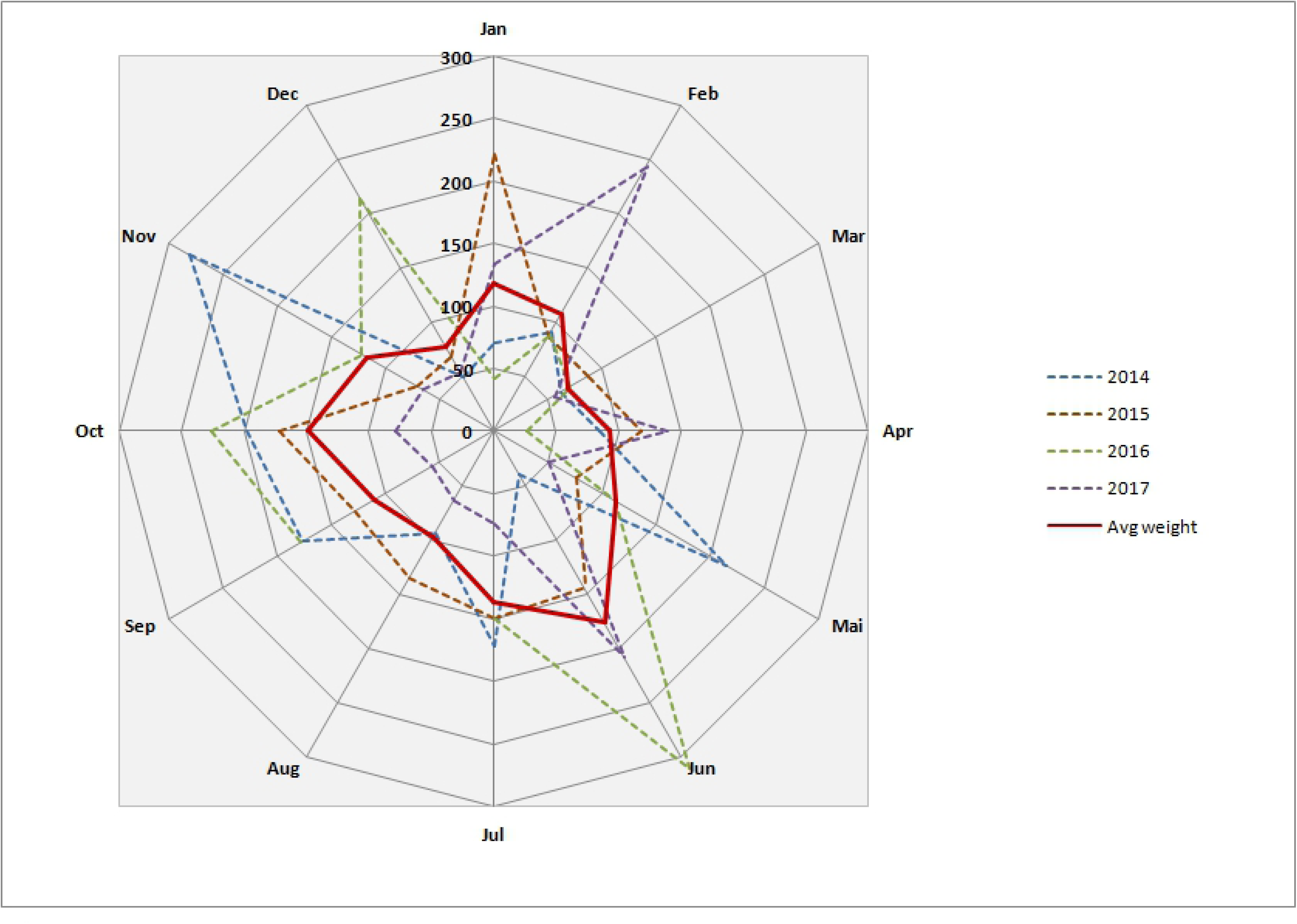
Weight (Kg) per month of dusky grouper *Epinephelus marginatus* from Copacabana small-scale fishery.

Statistical comparisons (i.e., the Kruskal-Wallis tests) showed that catches were stable among the years of sampling in Copacabana. The results of the statistical comparisons of 4 full years (a) (2014-2017, Table 1) and 6 partial years (b) are as follows:

**Table 1.**
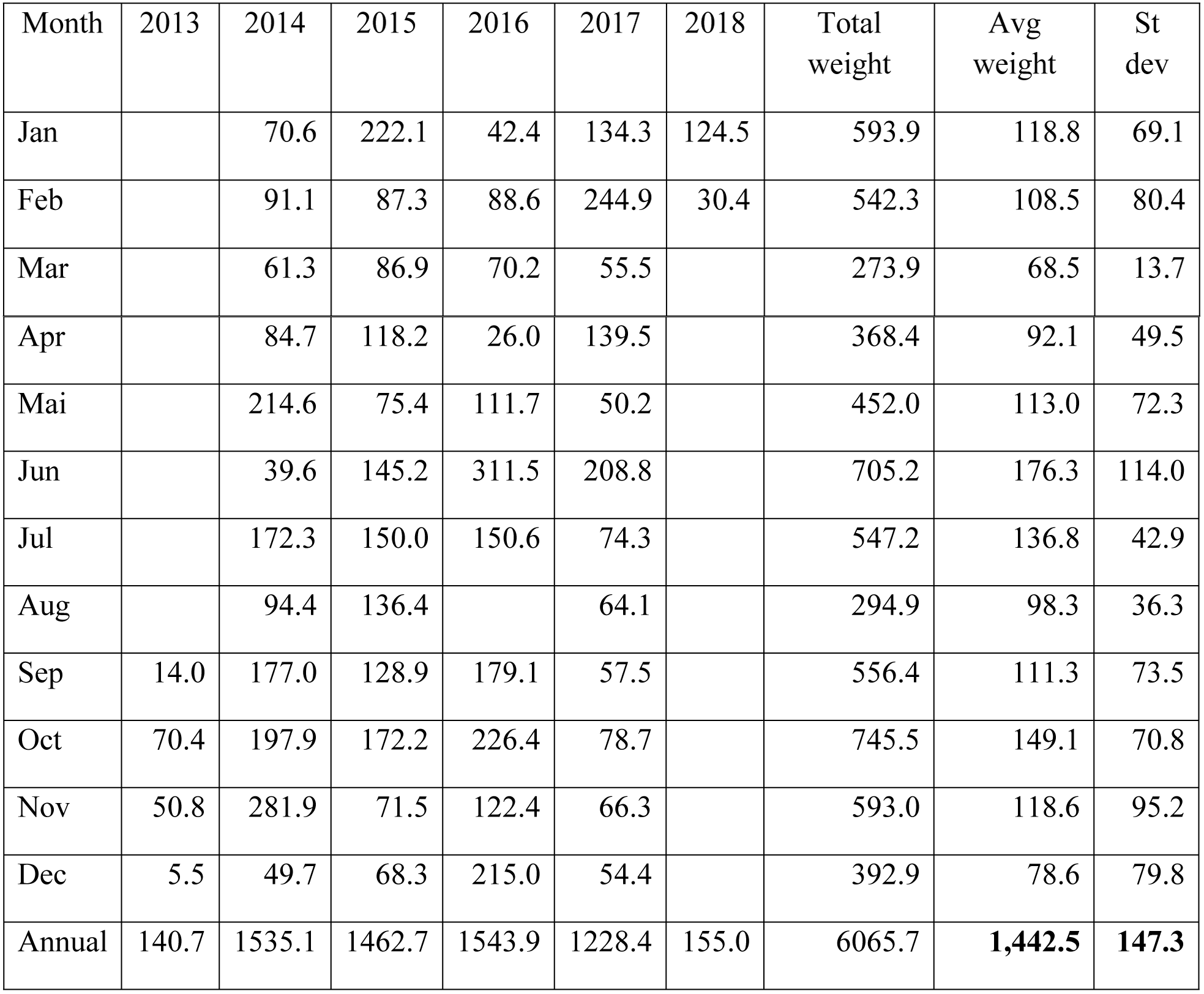
Kg per month: table with all years and total per year; averages and std^1^ of dusky grouper, *Epinephelus marginatus* at Copacabana, Rio, 2013-2019 (n=1,896 individuals) ^1^averages calculated for period of full years, 2014-2017.

a. The total number of observations was n=48; the sum of the ranks of the total number of observations was 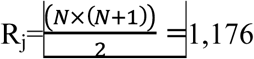 (S1 Table). The parameters are α=0.05 and df=4-1=3; the χ^2^ value is 7.81.
b. The total number of observations was n=54. The sum of the ranks of the total observations in the first version was 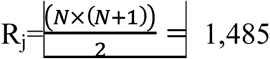 (S2 Table). The parameters were α=0.05 and df=6-1=5; the χ^2^ value was 11.07.

The critical H value (H_c_) was defined from the χ^2^ criterion.

In both cases, the calculated values of the Kruskal-Wallis statistics were lower than the H_c_. Thus, we concluded that there were no significant differences between grouper catches of different years.

The very productive months, whose productivity was calculated from the average weight of the dusky grouper, were June and July (i.e., the winter) and October (i.e., the spring).

### Weight-length of dusky groupers

The weight-length of groupers from the 6-year period (2013-2019) is shown in Fig 2. Most groupers fall between 50 and 70 cm. The figure was based on 1,563 individuals, and the curve is expressed by W=0.0043*L^2^-0.3122*L+6.9191, R^2^=0.925 (n=1,563; df =1,561)

**Fig.2.**
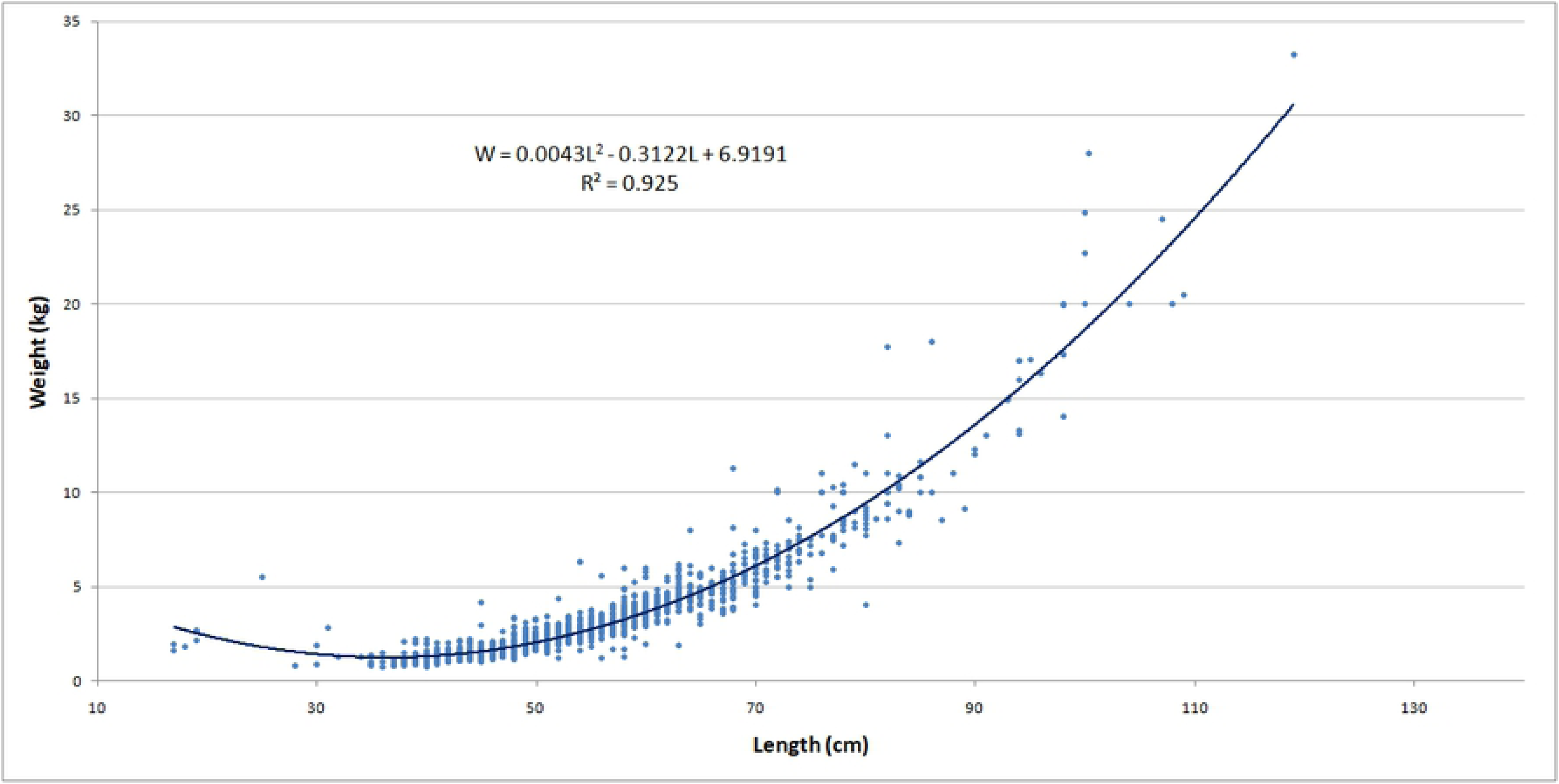
Weight-length (kg-cm) of dusky grouper *Epinephelus marginatus* from Copacabana small-scale fishery.

Fig 3 clearly demonstrates these results, with a peak between 50 and 60 cm, which corresponds to 37% and 31% of the groupers caught, respectively (n=1,544 individuals).

**Fig. 3.**
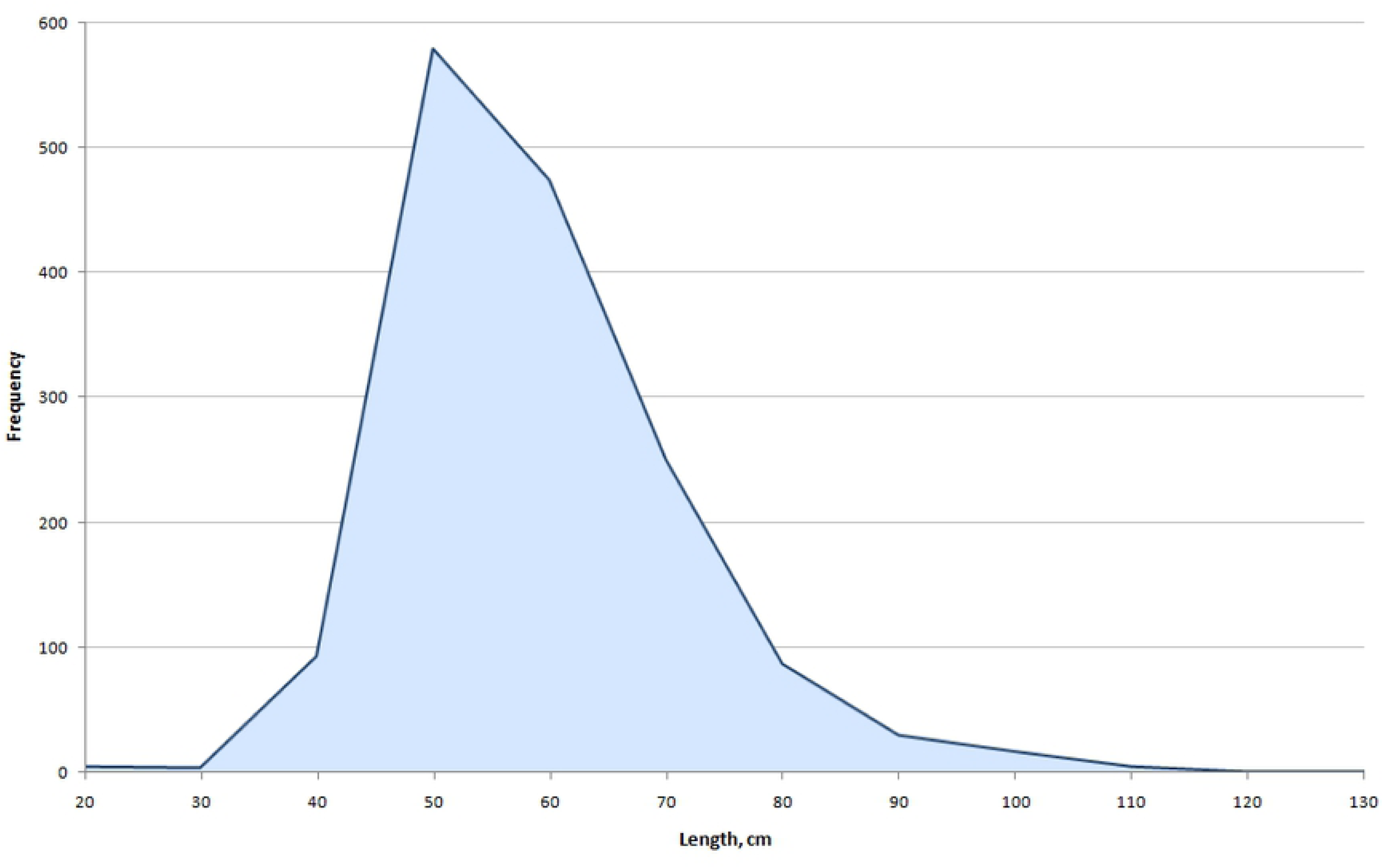
Length (cm) of dusky grouper *Epinephelus marginatus* from Copacabana small-scale fishery.

### Reproduction

From 515 days of sampling of the dusky grouper in Copacabana, we observed 1,969 individuals, of which only 1,83% had visible gonads with eggs. The mean volume of the gonads was 54.71 ml, and the fish had a mean weight of 5.96 kg and a mean length of 65.14cm (Table 2). Approximately 10% of the fish with visible eggs or gonads occurred in the spring (October and November) and a few occurred in the summer and autumn (January-March and April, respectively). The peak of the dusky grouper catches is concentrated in the winter, when the fish are probably not in their reproductive period.

**Table 2.** Month (summing up years 2013-2019): visible gonads, mean size and weight of fish with visible gonads. Including total number of individuals of dusky grouper, *Epinephelus marginatus*, sampled, total days sampling, total number of individuals with visible gonads

| Month | Days sampling | Number of individuals sampled | Fish with visible gonads | % fish with visible gonads | Mean gonads volume, ml | Mean Weight, kg | StDev Weight | Mean Lenth, cm | StDev Lenth |
| --- | --- | --- | --- | --- | --- | --- | --- | --- | --- |
| Jan | 49 | 191 | 2 | 1.05 | 29.5 | 5.00 | 0.99 | 63.00 | 1.41 |
| Feb | 41 | 148 | 5 | 3.38 | 93.0 | 7.35 | 6.42 | 68.80 | 15.45 |
| Mar | 32 | 102 | 1 | 0.98 | 150.0 | 7.20 | - | 78.00 | - |
| Apr | 28 | 104 | 4 | 3.85 | 25.0 | 7.61 | 3.15 | 64.75 | 27.83 |
| Mai | 38 | 141 |  | - |  |  |  |  |  |
| Jun | 57 | 254 |  | - |  |  |  |  |  |
| Jul | 58 | 199 |  | - |  |  |  |  |  |
| Aug | 36 | 97 | 1 | 1.03 | 20.0 | 7.61 | - | 77.00 | - |
| Sep | 49 | 187 |  | - |  |  |  |  |  |
| Oct | 60 | 228 | 9 | 3.95 | 12.1 | 5.40 | 3.28 | 62.33 | 12.58 |
| Nov | 38 | 183 | 11 | 6.01 | 76.2 | 5.16 | 4.24 | 62.18 | 15.72 |
| Dec | 29 | 135 | 3 | 2.22 | 30.0 | 5.81 | 1.75 | 72.00 | 5.00 |
| **Total** | **515** | **1969** | **36** | **1.83** | **54.71** | **5.96** | **3.79** | **65.14** | **14.89** |

### Fishing spots

Groupers were often caught in rocky shores, and some islands were of overwhelming importance for catching dusky groupers. Table 3 shows the importance of the islands in the Cagarras Archipelago (including the fishing in the vicinity of these islands, which fishers refer to as “ao largo das cagarras” and “laje da cagarra”), and the Rasa and Redonda Islands (which account for most of the biomass caught). From Cagarras, Redonda, and Rasa, groupers represent 42%, 13%, and 7% of the biomass caught, respectively; compared with Cagarras, Redonda and Rasa are relatively distant islands. Fig 4 graphically shows the biomass of groupers per fishing spot, and S1 Fig. shows a map with the fishing areas and fishing spots.

**Fig. 4.**
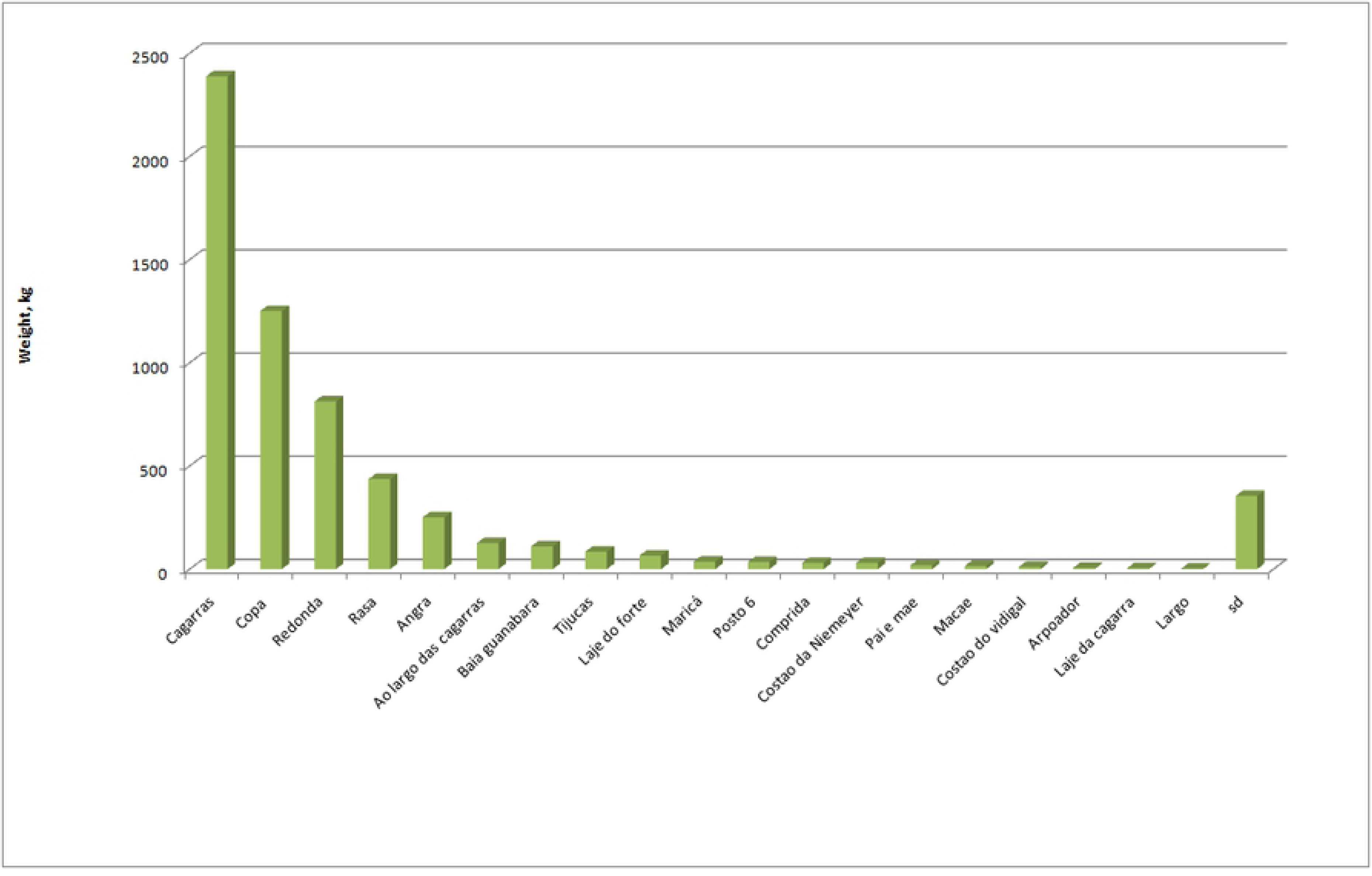
Fishing spots and production per spot (kg) of dusky grouper *Epinephelus marginatus* from Copacabana small-scale fishery.

We had the opportunity to include some data from recreational fishers of the adjacent Marimbás Club. Data from sixty-nine trips (in 2015 and 2016) were included, equaling 338,12 kg of dusky groupers caught at Cabo Frio in Northeast Rio de Janeiro state. The average weight of dusky grouper was 4.9 kg, slightly above the average weight of dusky grouper from the main spots used in Copacabana (Cagarras, Rasa and Redonda, where the approximate average weight of dusky grouper was 3.3 kg). Distant locations, such as Angra, Maricá and Macaé, showed the highest average weight per fishing spot (Table 4).

**Table 4.** Fishing spots and kg per fishing spot at the fishery of Copacabana where dusky grouper *Epinephelus marginatus* was caught.

| LOCAL | Weight (Sum),<br>kg | individuals | Avg Weight of ind. | StDev-Weight |
| --- | --- | --- | --- | --- |
| cagarras | 2387.80 | 724 | 3.32 | 3.0 |
| copa | 1249.10 | 439 | 2.85 | 1.8 |
| redonda | 812.09 | 244 | 3.33 | 2.7 |
| rasa | 435.95 | 135 | 3.23 | 2.4 |
| angra | 251.00 | 53 | 4.74 | 4.7 |
| ao largo das<br>cagarras | 126.24 | 45 | 2.81 | 2.4 |
| baia guanabara | 110.22 | 43 | 2.56 | 1.3 |
| laje do forte | 65.25 | 26 | 2.51 | 1.7 |
| tijucas | 84.21 | 23 | 3.66 | 3.7 |
| costao da<br>Niemeyer | 30.28 | 14 | 2.16 | 0.8 |
| posto 6 | 34.39 | 14 | 2.46 | 1.4 |
| costao do vidigal | 10.70 | 6 | 1.78 | 0.5 |
| comprida | 30.40 | 5 | 6.08 | 6.2 |
| maricá | 36.30 | 3 | 12.10 | 7.0 |
| laje da cagarra | 4.64 | 2 | 2.32 | 1.3 |
| pai e mae | 18.30 | 2 | 9.15 | 5.9 |
| arpoador | 5.53 | 1 | 5.53 | - |
| costão | 1.40 | 1 | 1.40 | - |
| largo | 2.79 | 1 | 2.79 | - |
| macae | 14.90 | 1 | 14.90 | - |
| sd | 354.21 | 118 | 3.00 | 2.6 |

### Gear used

In Copacabana, spearfishing (by snorkeling and probably some diving) is the most common method to fish dusky grouper (Fig 5).

**Fig. 5.**
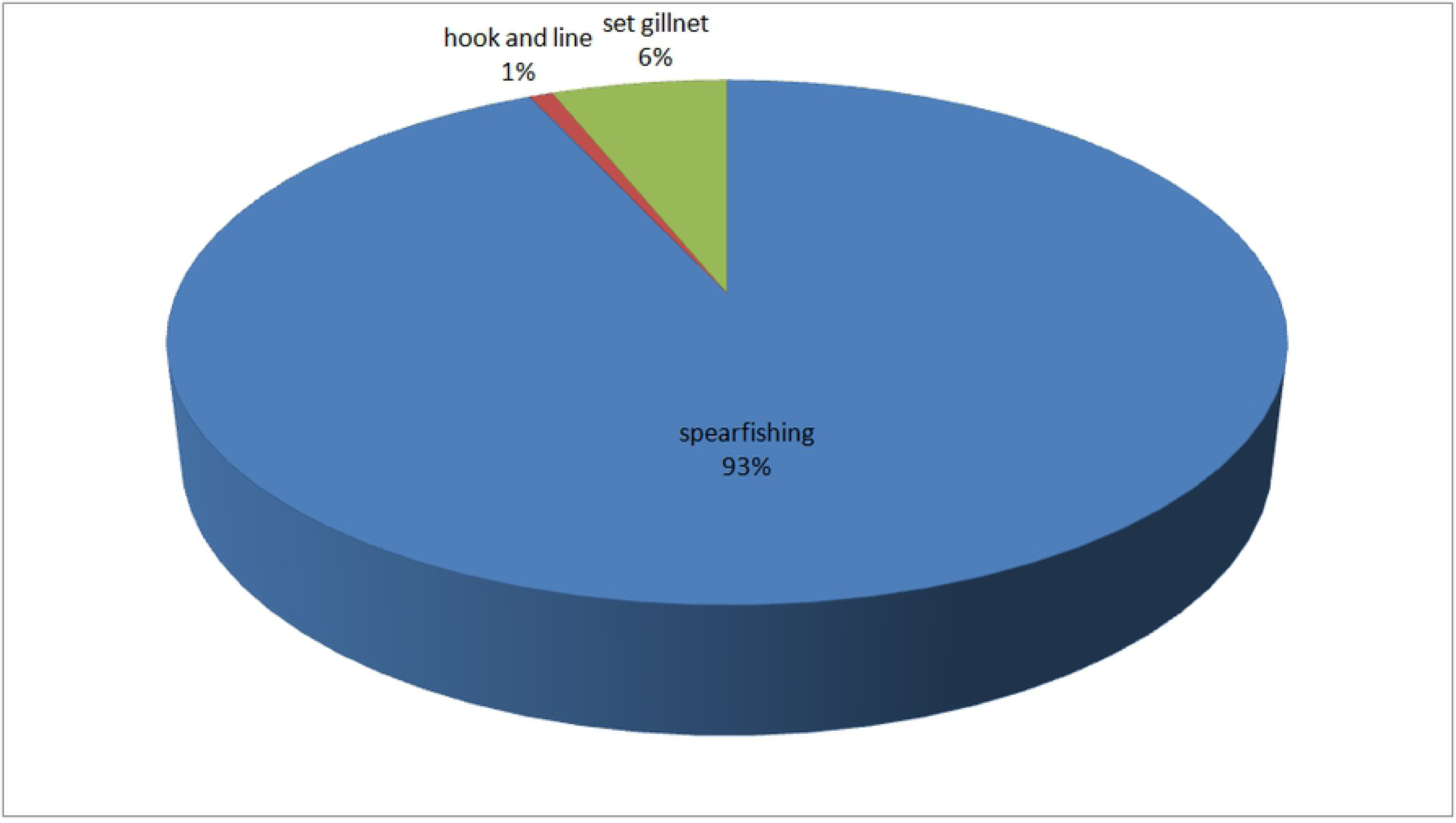
Gear used to catch groupers at Copacabana.

### Prices

Data on prices are given in Figs 6 and 7 with details in S4 Table and S1 and S2 Figs. The dusky grouper prices increased 2.774 reals per year. Considering that the average price was approximately 34 reals/kg in 2016, the price increase corresponds to an increase of approximately 8%, which is similar to the figure of inflation in Brazil, which was 6,29% in 2016, 2,95% in 2017, and 3,75% in 2018 (https://g1.globo.com/economia; http://agenciabrasil.ebc.com.br).

**Fig. 6.**
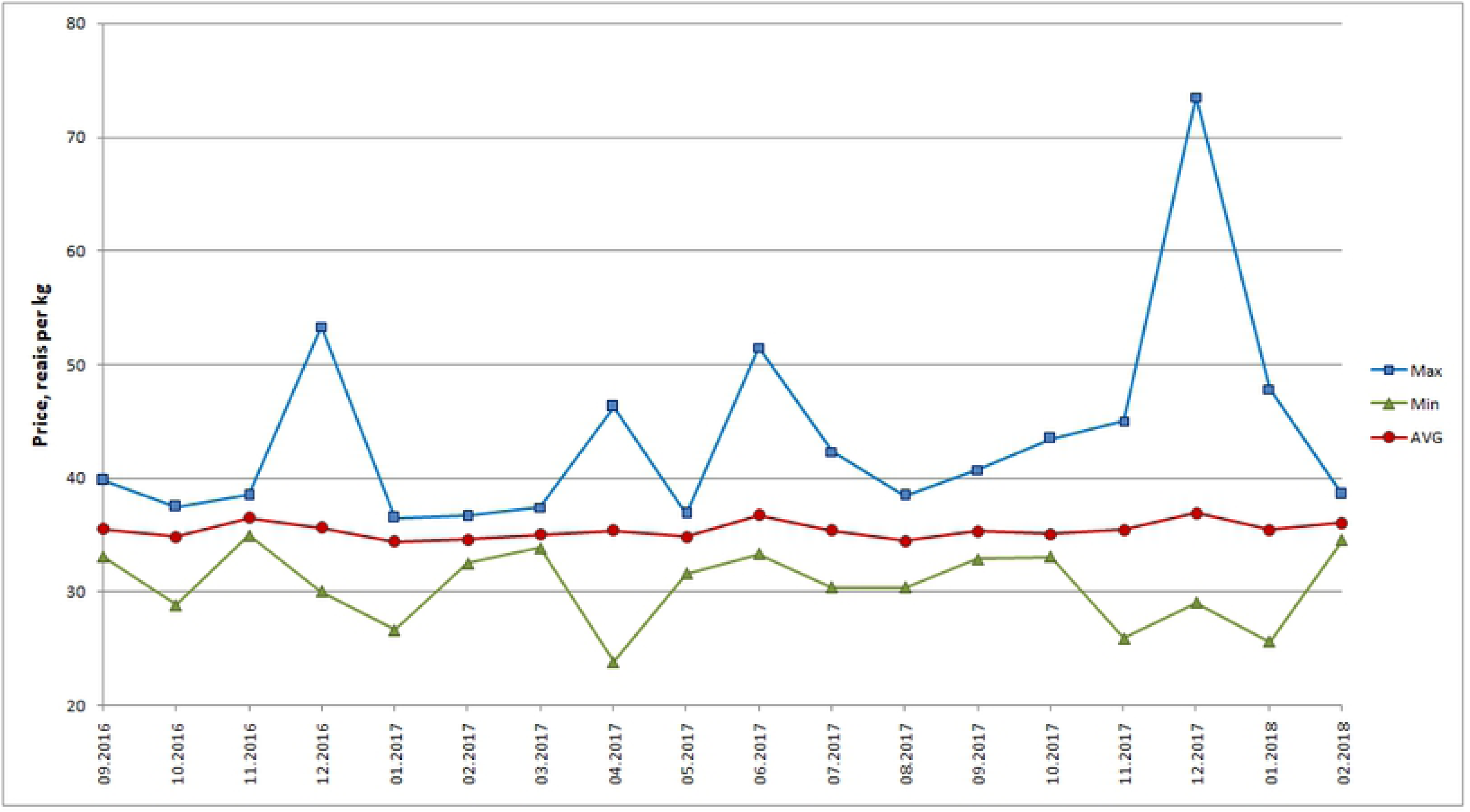
Prices (Reais, R$) of dusky grouper *Epinephelus marginatus* from Copacabana small-scale fishery.

**Fig. 7.**
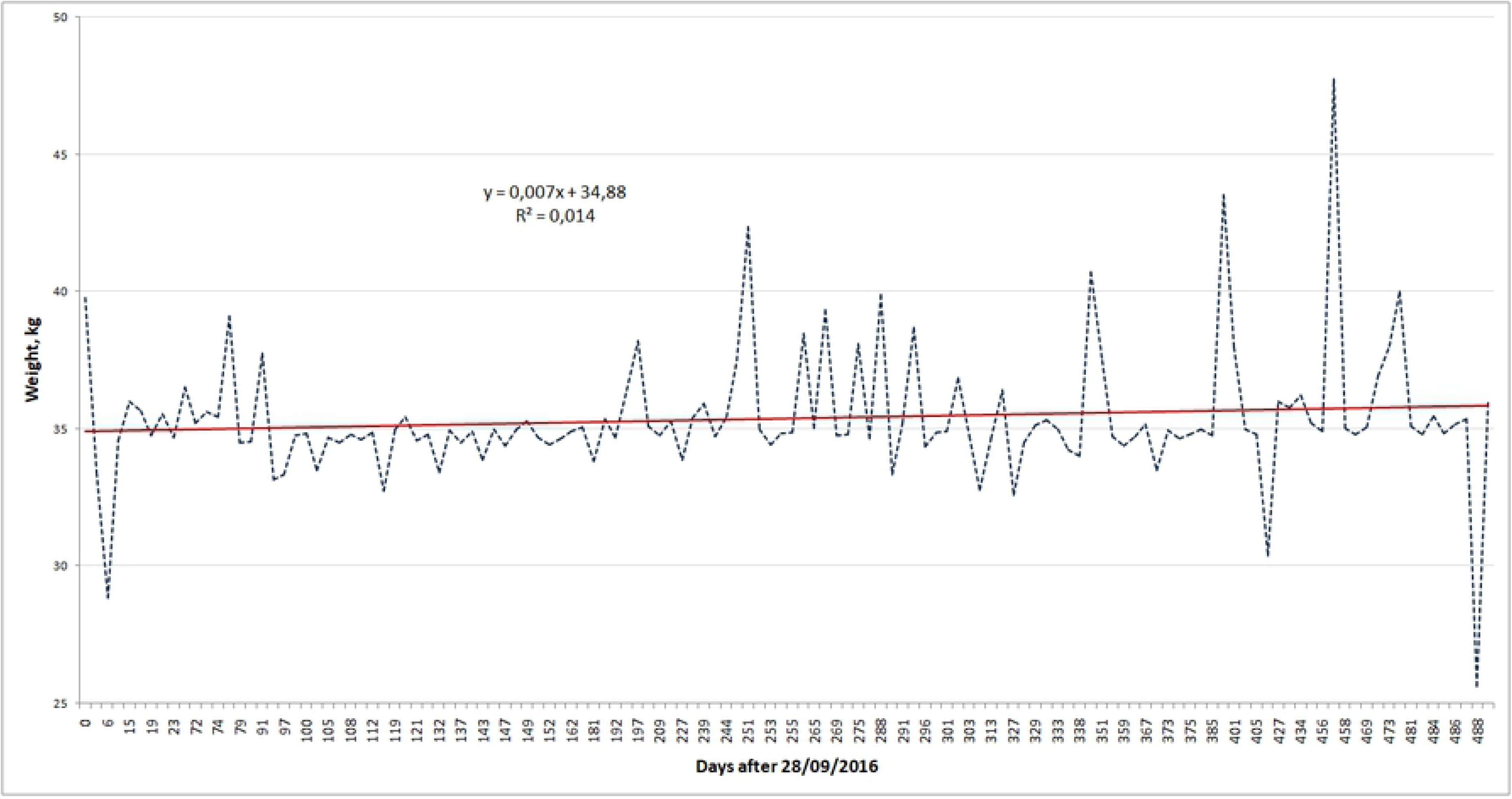
Average monthly price of dusky grouper *Epinephelus marginatus* for different years (2016-2019) from the small-scale fishery of Copacabana. Price Regression analysis showed that the average price of grouper (Reais, R$, per kg) increases by about R$ 2.774 per year

We compared samples with the data on the grouper prices for three adjacent years, from 2016 to 2018. Note that sample sizes for different years are quite different. We have 71 price values for the period from 28.09.2016 to 30.12.2016, 264 price values for the period from 03.01.2017 to 30.12.2017, and 50 price values for the period from 09.01.2018 to 10.02.2018.

S4 Fig shows that the normalized histograms of the price distributions look similar. However, the data for 2016 and 2018 have wider and lower peaks shifted toward larger price values. Considering the aforementioned, for our Kruskal-Wallis analysis, we used prices averaged per day and divided the 2017 data into quarters. Thus, we obtained 5 independent samples with a similar number of observations corresponding to time periods of approximately the same length. The total number of observations is n=128.

The sum of the ranks of the total observations is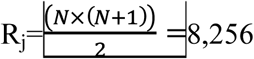.

The distribution of the number of observations and the rank sums for each sample is given in S3 Table.

H_c_ was defined from the χ^2^ criterion for the parameters α=0,05 and df=5-1=4. The χ^2^ value was 9.49.

The calculated value of the Kruskal-Wallis statistic was lower than the critical value (H=8.42<H_c_=9.49). Thus, we concluded that there were no significant differences between price samples for different years.

## Discussion

Compared with the results of a 2016 study on dusky grouper in Copacabana, we had double the sample size and a more significant regression on the weight-length ratio (R^2^=0.83, df=791 in the 2016 study; R^2^=0.92, df=1,561). Increasing the sample size also allowed us to conclude that the catches in Copacabana included larger dusky groupers than the previous study found. Diving continues to be the main method to catch dusky grouper, and the fishing spots remain the same near the islands of Cagarras, Redonda and Rasa.

### Bottleneck

Considering that the dusky grouper is an endangered species, analyzing the catches in Copacabana relative to those in previous studies on its genetics from this area is worthwhile. The population of groupers from the sites in Copacabana and Paraty, where the original catch data used to perform genetic analyses was collected, showed a significant bottleneck, with a population of 663 individuals [23].

Small populations show genetic effects such as bottlenecks and drifts (i.e., random effects) as well as inbreeding depression (i.e., directional effects); bottlenecks indicate that the population size declined [24]. When there is a sudden reduction in population size, the average heterozygosity per locus is expected to decline [25]. Survivors carry a sample of the genetic variance from the prebottleneck population; the minimum effective population sizes to minimize the loss of variance and fitness were suggested to be 50 for short-lived individuals and 500 for long-lived individuals [24].

The stock structure of most grouper species is not well understood [26]. However, data from the Priolli et al. [23] study showed a bottleneck in the population of dusky groupers from the sites in Copacabana and Rio-Paraty, in southern Rio de Janeiro. Considering our results and the comparison of yearly catch data, we can conclude that there is a sustainable fishery of dusky grouper from Copacabana, Posto 6, that has the capability of maintaining a stable population of dusky groupers. Moreover, it is important to observe that studies of dusky grouper populations in Malta have shown decreasing catches, with a mean length of 52 cm (n=84) and an average weight of 3 kg (n=41) [27]. Our larger sample in Copacabana showed an average weight of 3,20 kg (n=1896; std=2,74) and length of 54,47 cm (n=1544; std=12,11).

### Prices

Our price samples comprised 128 individuals of dusky grouper from throughout the year, and showed a non-normal distribution. There were no significant differences among the years in the price of dusky grouper in Copacabana, Rio de Janeiro.

Dusky grouper prices reflect the lack of variation in grouper catches over the years and reinforce the stable availability of groupers in these years. In the Gulf of Mexico and the South Atlantic areas, the prices of fishes of the snapper-grouper complex were analyzed using SIDS (synthetic inverse demand system). While fishing probably responds to fish price incentives, catches are not influenced by the probable random perturbation in the monthly fish price. In a study about fish prices in India, Sathiadhas and Kumar [28] emphasized the important points concerning marine fishery prices: marine fish prices are very uncertain due to the unpredictability of production, fish are highly perishable, landing points and species are diverse, there are spatial and temporal variations, and there are disequilibria of supply and demand, among others. When they are stressed, fish fluctuate far more than agricultural products, and the short-run supply is highly inelastic: an increase in catches will lower prices, and a low catch will increase prices.

Fish show an inverse demand system: the prices reflect the quantities of fish in this inelastic system where the producers tend to be those collecting the money (there is no fish processing before the catches are landed in the local market) [29,30]. Gates [31] showed how important fish size is for determining prices. In the case of groupers, the price is given per kg; of course, this differs between fish species and groupers are considered a noble fish, with a high price in the market of Copacabana [14]. As shown by optimal foraging models, noble species tend to be less bony and, thus, require less manipulation time [21,32]. Thus, from these results, we may assume that the stable average prices are due to stable catches throughout the years. This is a very important result, since fishers have felt repressed from their fishing activities, especially for noble species. Recent legislation (Portaria Ministerail 41, July 27, 2018) related to the fishing of dusky grouper intends to provide management but without any participation of fishers.

As with marine fisheries from other developed countries, harvesting and marketing of fish create employment opportunities [28]. This applies to the fishery of Copacabana, where fishers have depended on the fishery for a living for several years [14,15]. Knowing that the grouper fishery has been sustainable is very important, especially because it is a noble fish with high market prices [14]. Unfortunately, another legislation, the Decree 8722 from May 11, 2016, has been an obstacle to both small-scale fishers and researchers since it inhibits fishing and collaborative processes between fishers and researchers.

### The future: systems of knowledge-citizen science and local ecological knowledge and their importance for data-poor small-scale fisheries

There are many instances in which local ecological knowledge has been useful. An example of the different knowledge systems available for a cosmopolitan species is found in the studies from Australia and Brazil on *Pomatomus saltatrix* (bluefish) [20, 33-35]; in this very interesting example, the local knowledge of bluefish in Australia and in Brazil yields enhanced data on its migration and reproduction, among other characteristics. In the eastern Australian state of Queensland, Brodie et al. [33] conducted a tag-recapture CS study, and the spatiotemporal movements of *P. saltatrix* were recorded; thus, important information for the management of bluefish became available. In Brazil, [34,35] studies on the LEK of the same species (*P. saltatrix*) were performed by comparing the LEK regarding feeding, habitat and migratory movement direction of fishers from Buzios Island (SE Brazil) to the aboriginal fishers of North Stradbroke Island in Moreton Bay (Queensland) (*Quandamooka* in the aboriginal language); both sets of LEK were also compared to the information from the literature, showing an astonishing correspondence. Reproduction data and other important ecological and biological types of information were also obtained in the studies of LEK and CS. Brodie et al. [33] observed that some species lack information regarding catch records because several factors prohibit them from being targeted; however, recreational fishing programs are similar to programs of citizen science in which members of the public participate in data collection, e.g., by tagging and releasing fish.

Postuma and Gasalla [36] presented results from fishers’ information (i.e., LEK) about squid, which they used to cross-validate their analysis and provide important information about the decrease in the concentration of squid in SE Brazil, and they gave the environmental conditions of the areas where the best squid catches were known to occur in this region. Mapping is also an important aspect in which LEK is very helpful and the knowledge of geographical information related to fishing spots has been shown to be helpful in some studies. For example, Léopold et al. [37] mapped mangrove and coral reef finfish and invertebrate fisheries in New Caledonia (in the southwest Pacific). In Brazil, several species were mapped using GPS with the help of local fishers [38]; in particular, in SE Brazil, important target species were mapped through this method, including catfish, mackerel, snapper, croaker, sand drum, bluefish, cutlass fish, grouper, weakfish, snook, mullet, bluerunner, shrimp and squid [15].

Information on food chain and feeding habits was also provided, with LEK as a source among other relevant biological information [39-43]. This is especially important where there is nonexistent information, or a few sources of information, on fish, such as on the coast of Brazil [18]: there are certain species for which the only source of knowledge is LEK, such as *Rhinobatos percellens, Sphoeroides dorsalis*, and *Dasyatis guttata*. It is important to note that information is lacking for important target species and consumed species from both the coastal and continental waters of Brazil [9,10]. Another area in which the fishers can help is the monitoring of contaminants: as the study by Silvano and Begossi [44] shows, observations of bioaccumulation, such as the mercury content in fish muscle, can be made by fishers and can be very valuable.

Other information obtained through LEK provides insights for modeling and predicting the distributions of species. This was shown in a study by Lopes [45] for the coast of Brazil using Bayesian hierarchical spatial models and oceanographic variables, with which the author was able to predict the distribution of grouper, *Epinephelus marginatus*; they used data from the literature and from fishers, and the results showed a concordance between the models that temperature predicts the distribution of this species and the reliability of the information from the fishers. Duplisea [46], studying redfish (*Sebastes* spp.) catches, concluded after interviews with fishers, that scientific reports underestimated catches, including those of small fish, which led to a reinterpretation of the abundance of stocks. Fishers are also shown to provide helpful information regarding the temporal abundance of fish and changes therein, based on CPUE [47] and regarding different fishers’ perceptions concerning MPAs. Other temporal changes were also detected by Lima et al. [48] in the SE Atlantic Brazilian areas, such as a decrease in biomass of the fish caught and a higher abundance of relatively small fish.nats

### LEK in freshwater fisheries

In a recent study by Nunes et al. [49], LEK of migratory behaviors and other ecological information from seven Amazonian fishes along a 550 km stretch of the Tapajos River was studied through interviews with 270 fishers, who also provided information on the behavior of this fish along the poorly known tributary rivers of the Amazon; this information was of great importance considering dam building in these areas. Changes in fish abundance have also been identified through LEK: Hallwass et al. [50] demonstrated this in the lower portion of the Tocantins rivers by acquiring information from 300 fishers in 9 villages and 601 fish landings, and concluding that, after impoundment of the river due to the Tucuruí dam, an important local species (jaraqui, *Semaprochilodus brama*) had become locally extinct, and there had been changes in the composition of fish catches and decreases in fish production.

In the Mekong River in Asia, studies by Valbo-Jørgensen and Poulsen [51] involving 355 expert fishermen along 2,400 km of the Mekong mainstream were used to develop migration maps and obtain spawning information for 50 fish species.

### Citizen science: examples

Citizen science (CS) (i.e., public scientific research, [52]) refers to the knowledge that local people have about something or might refer to their collaboration to some ongoing research; it has been applied to ecological and biological knowledge and it can fulfill the research demands from scientific points of view and incorporate rigorous research tools, as demonstrated by the many examples given in this study by Cigliano et al. [52]. Bailey et al. [53] suggest that a positive relationship between fishery scientists and lay people could bring positive results toward integrative research as a “social activity”.

Other examples are given by Fairclough et al. [54] concerning citizen science projects being a cost-effective form of data collection because of the opportunity to obtain volunteers and increase the data sets and data coverage, among other effects. Spatiotemporal information was obtained using citizen science, and researchers were able to train and use focal groups to study inshore fisheries in Denmark [55]. The study of alien species [56] is another important aspect for fisheries management where CS is able to help in gathering ecological data.

### A gradient for dusky grouper: local knowledge and citizen science

Some research includes studies in which both local knowledge and citizen science are at work: examples include research in which fishers are trained to look for mature gonads in fish [19]. Where there are no data on fish, fishers are extremely helpful since they can gather data, thereby increasing the reliability of sampling and results. Projects in Brazil and other studies were performed in this way, especially with grouper (*Epinephelus* spp.) [57,58]. These are studies in which both local knowledge and citizen science are involved; the local knowledge comes from fishers who provide their own knowledge (such as the reproductive period or knowledge on the aggregation of species), but the fishers also participate in a training program within the research project. Thus, the two forms of knowledge are involved in a “dialogue between different forms of knowledge”, for which Ruddle [59] provides several examples and details.

Both methods and concepts, i.e., local knowledge and citizen science, can be helpful not just for data-poor countries: management in the North Sea shows several weaknesses, indicating that there is a need to integrate citizens because fishers can communicate as both citizens and knowledgeable experts with defined interests at stake. Indonesia has high average shark landings, but with very little local information [60]; this is another study conducted in a particularly data-poor region where sharks are primarily targeted for their fins with the help of shark fishers.

These results are very important, especially considering that dusky grouper, *Epinephelus marginatus*, is a flagship species for conservation that reaches large sizes and lives up to 60 years, with many MPAs (marine protected areas) established in Mediterranean waters [61]. It is a very important food source for coastal communities that have livelihoods dependent on fishing [9,18], and it is an ecological and cultural keystone species [10].

## Conclusions

Our study shows that, despite the vulnerable features of dusky grouper, the small-scale fisheries in Copacabana have been performing well, since grouper catches have been stable over time. In data-poor fisheries such as those in the coastal communities of Brazil, an integration of science and citizen science (including local knowledge) can turn viable scientific conclusions into management practices.

The integration between the different types of knowledge has been put into practice by the FAO [62], with a technical paper about the Latin American fisheries providing a toolkit for an ecosystem-based management approach. Several years ago, in 1981, Johannes [63] explained how important fishers could be for managing resources and data-poor fisheries; approximately 20 years have passed since the publication of classical papers on “ignoring fishers’ knowledge and missing the boat”, among other topics [64,65].

As shown in this study, the dialog between other forms of knowledge and scientific research can be used in the following ways to help in fisheries management, especially in data-poor areas:

1. Increasing the possibility of monitoring aquatic species.
2. Increase sample size.
3. Organizing systematic data collection with the help of expert fishers or citizens.
4. Diminishing research costs in terms of sampling size (i.e., by making increasing sampling possible) and monetary expenses.
5. Sharing knowledge with other nonscientific groups, such as expert fishers’ and the public.
6. Learning from other nonscientists who show empirically-based knowledge, such as expert fishers and the public.
7. Increasing the possibility of knowing important factors, such as the period of fish reproduction and reproductive aggregations, which are key factors for fisheries management (and are unknown in several data-poor countries).
8. Increasing the knowledge of target species, such as dusky grouper.
9. Increasing the knowledge of vulnerable species, such as dusky grouper.

There are examples from Mexico that should be followed, showing how important the participation of fishers is in the management of groupers [66]. Other examples for Latin American and the Caribbean are found in Salas et al. [2].

## Acknowledgments

I am thankful to FAPESP (#14/16939-7) and CNPq (301592/2017-9).

This research is approved and signed by B. R. Martins dos Santos, Comitê de Ética, Universidade Santa Cecília, number 1.747.889 on September 27, 2016 (Plataforma Brasil). It is approved under number 53824 at SISBIO and registered under number AB53669 at SISGEN, MMA (Ministério do Meio Ambiente, Brasil). We are grateful for fishers Antonio, Elenilson (“Jaguriçá”) and Raul; Ana was helpful with data from the Marimbás Club, Copacabana.

